# A comprehensive identification and annotation of mobile LINE-1 retrotransposons across human reference and personal genomes

**DOI:** 10.1101/2025.02.17.638711

**Authors:** Adriano Ferrasa, Roberto H. Herai

## Abstract

LINE-1 (L1) is a type of mobile genetic element (retrotransposon) that can self-copy into distinct parts of its host genome by dynamically shaping its content. In humans, misregulation of these elements have been associated as the potential cause of distinct types of pathologies and neurodevelopmental disorders, including cancer, Rett syndrome and Aicardi Goutierrez disease. However, the precise annotation of these 6 kb long nucleotide sequence is a challenging task due to the high similarity between their internal regions (coding and non-coding) for distinct L1 families. Current L1 reference annotations such as RepeatMasker still lack sequence details and also provide confounding annotated information, that includes thousands of incomplete L1 sequences annotated as representative full-length, and missing information on their regulatory and coding regions (5’UTR, ORF1, intron, ORF2 and 3’UTR). Additionally, current L1 resources also provide annotations based on consensus reference genomes and does not include individual genome information at allele-level and nucleotide-level resolution. Here, we present L1Farm as a complementary databank and annotation resource that contains a comprehensive analysis and annotation of L1 subfamilies identified in the current human reference genome (build Hg38) and in two individual Asian and Caucasian genomes at allele-level and nucleotide-level resolution. L1 elements sequences were also annotated with genomic *loci* and ranked according to their full-length content and similarity.

## Introduction

Long Interspersed Nuclear Element type 1 (LINE-1 or L1) are replicative repetitive elements and make up a significant portion of 21% of human genome [1]. They have been evolving during at least 170 million years of mammalian adaptative radiation and represent an important mechanism of genomic evolution by dynamically reshaping its content [2–4].

As a type of mobile genetic element, L1 retrotransposons are able to self-copying into distinct parts of the host genome (retrotransposition activity) [5], and their activity varies according to the family they are classified, that is based on their nucleotide content [6]. Currently, L1 subfamilies are classified according to their age, with the 2 youngest known as L1HS and L1PA2. Along with the subfamily variations, the copies of each L1 can accumulate mutations at a neutral rate. Thus, based on their nucleotide sequence content, older L1 subfamilies are more divergent than younger families [4,5]. In humans, only copies from most recently evolved L1 subfamily, L1HS, are capable to retrotranspose [7–10] and their expression is modulated by several host cellular proteins, such as TREX1 (*Three Prime Repair Exonuclease 1*), p53 *(Tumor Protein P53*), RUNX3 (*RUNX Family Transcription Factor 3*), SOX2 (*SRY-Box 2*) and SP1 (*Sp1 Transcription Factor*) [11–15]. In certain diseases or disorders, such as cancer [16], Aicardi-Goutières Syndrome [11] and Rett syndrome [17], the expression of these proteins is unbalanced, suggesting that, although it has to be proven, the activity of L1 could be unstable and influence the pathogenic phenotype in each specific condition (reviewed in [16]). In other diseases, the genomic alterations driven by L1 retrotransposition activity has already been associated to psychiatric and neurodevelopmental conditions, such as schizophrenia and bipolar disorder [17–19].

Dispersed within the human genome, each copy of a full-length L1 element has ∼6 kb in length and encodes a 5’ UTR (untranslated region) containing an internal RNA polymerase II sense strand promoter [20] and a weak promoter on the antisense strand, a 3’ UTR that ends in a poly (A) tail, two main open reading frames (ORF), the ORF1 and ORF2 [21], and an inter-ORF spacer (hereafter intron) (Fig. 1). There is also a 216 nt antisense ORF0, locate at L1 5’ UTR locus that might be also involved in retrotransposition activity, but further investigations are required to demonstrate its role in L1 activity [22]. The ORF1 encodes a 40 kDa RNA binding protein (ORF1p) that has a nucleic acid chaperone activity [23,24], while ORF2 encodes a 150 kDa protein (ORF2p) with endonuclease and reverse transcriptase activities [25]. To be inserted as a novel copy within the host genome, the entire L1 element is transcribed as a polycistronic RNA, which is processed within the nucleus and then transported to the cytosol. Later on, this molecule is translated by the ribosomes to produce the proteins ORF1p and ORF2p. Together, these two proteins are associated with their encoding mRNA *in cis* to form a complex of ribonucleoprotein particle (RNP) [26,27] that are then transported back to the nucleus to begin the reverse transcription of the RNA. Then, the endonuclease of the ORF2p domain cleaves the genomic DNA by exposing a 3’ -OH which is followed by the removal of the L1 mRNA from the intermediate DNA:RNA hybrid, and the second strand DNA synthesis generates a new L1 insertion. This mechanism, called target-site primed reverse transcription (TPRT), generates a new L1 copy that is usually flanked by a sequence of 7 to 20 bp target-site duplication [28]. About 40% of L1 *de novo* insertions are identical to the initial L1 sequence and are capable of generating another retrotransposition event [29].

**Fig. 1.**
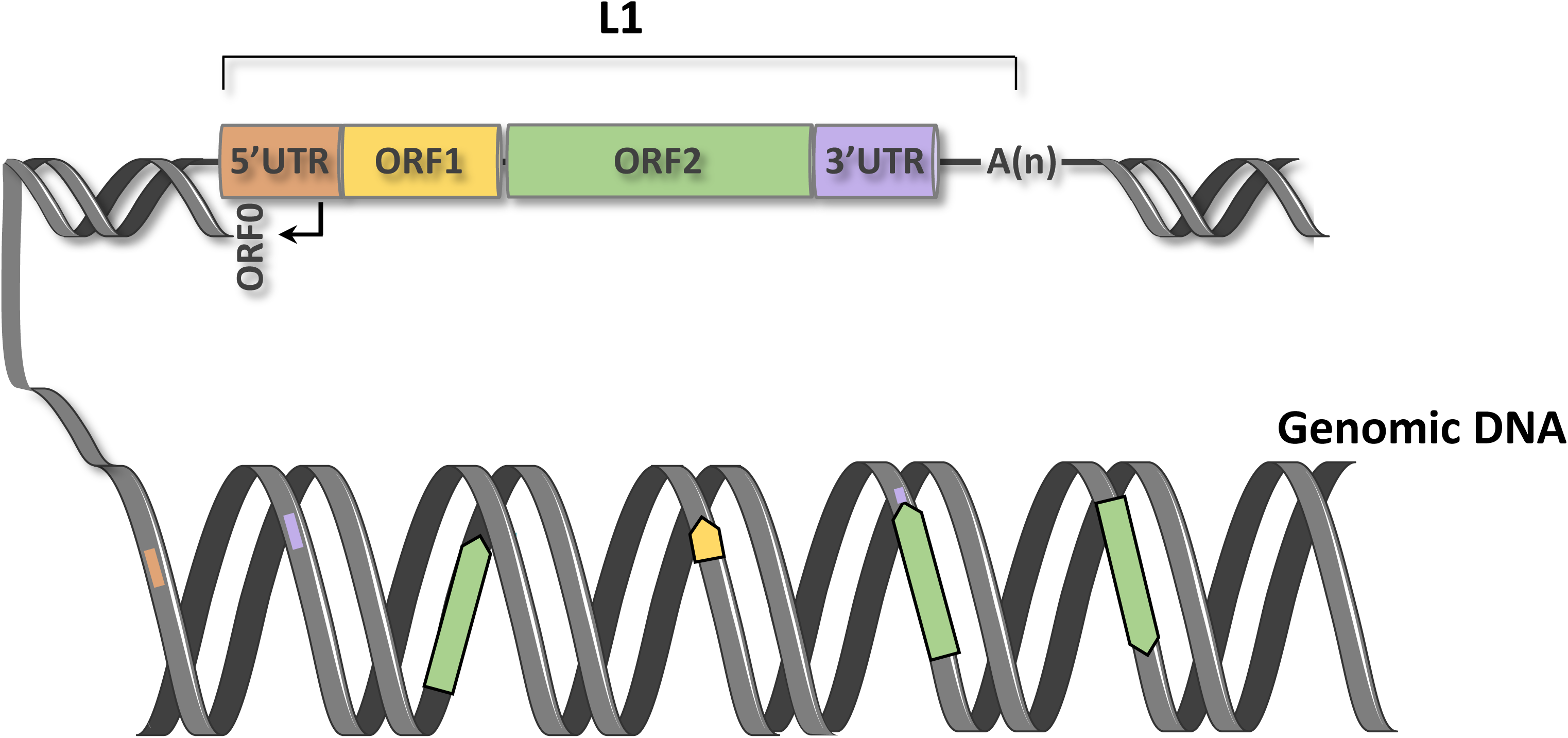
Schematic representation of a LINE-1 (L1) element. A L1 element consists of the following regions: a regulatory 5’UTR (shade orange line), a coding ORF1 (orange box), a coding ORF2 (green box) and a regulatory 3’UTR (shade blue). Between ORF1 and ORF2 open reading frames there is an intron region (black line). The 3’ UTR ends with a polyadenylation poly (A) tail (indicated as A(n)). L1 is spread throughout the genomic DNA and its insertions can be intact (full length integrations) or partial (entire sub-regions or a combination of them within random genomic *loci*).

Currently, there are several publicly available annotations of distinct L1 families and their corresponding genomic *loci*, such as RepBase [30], Dfam [31] and the most frequently used, RepeatMasker [32]. To detect and annotate the sequences within a target genome, including L1 elements, RepeatMasker uses a homology-based approach. It reports an annotation of distinct subfamilies of L1 elements across the entire human reference genome (Fig. 2A), including the most active retrotransposon L1HS subfamily element (1620 *loci* (copies) in the genome) and L1PA16 as the most repetitive in humans (13883 *loci* (copies) in the genome). However, an in-deep analysis of RepeatMasker to verify the subtype and length (length varying 250bp) of the annotated L1 elements (Fig. 2B) exposed some limitations, including several genomic *loci* having sequences shorter than 250 bp annotated as L1HS (a short 31 bp-length genomic region is annotated as L1HS, Fig. 2C). These short annotated sequences can lead to misinterpretation when analyzing the copies of ORF1’s present in L1 elements, since only some specific elements longer than ∼5 kb in length are able to transcribe and express as a RNA molecule with potential biological relevance [33]. The L1 ORFs, in addition to participating in the retrotransposition activity, can mobilize, *in trans*, thousands of processed pseudogenes as well as non-autonomous elements Alu and SVA (composite SINE/VNTR/Alu) [34]. Moreover, details regarding the L1 ORFs as well as the other L1 regions and full-length L1s are not provided through the RepeatMasker annotations. Furthermore, several annotated subfamilies of L1 sequences have overlapping annotations with at least 2 distinct molecules, not restricted to L1s [32]. Annotations available for L1 elements in RepeatMasker databank are also reported through the current assembly of the human reference genome (build Hg38), which is formed by large contigs and scaffolds, but several parts are still incomplete, including the telomeric and centromeric regions [35]. Moreover, the genome is represented as an haploid consensus sequence and originated from the genome sequence of multiple individuals [36]. Therefore, the availability of a complementary resource to repetitive databanks of L1 retrotransposons that better characterize and annotate L1 elements with detailed information, including similarity-based sequence annotations is required. It can improve *in silico*, *in vitro* and *in vivo* studies to improve our understanding of the role of retrotransposons in human genome in the definition of cellular and molecular phenotypes.

**Fig. 2.**
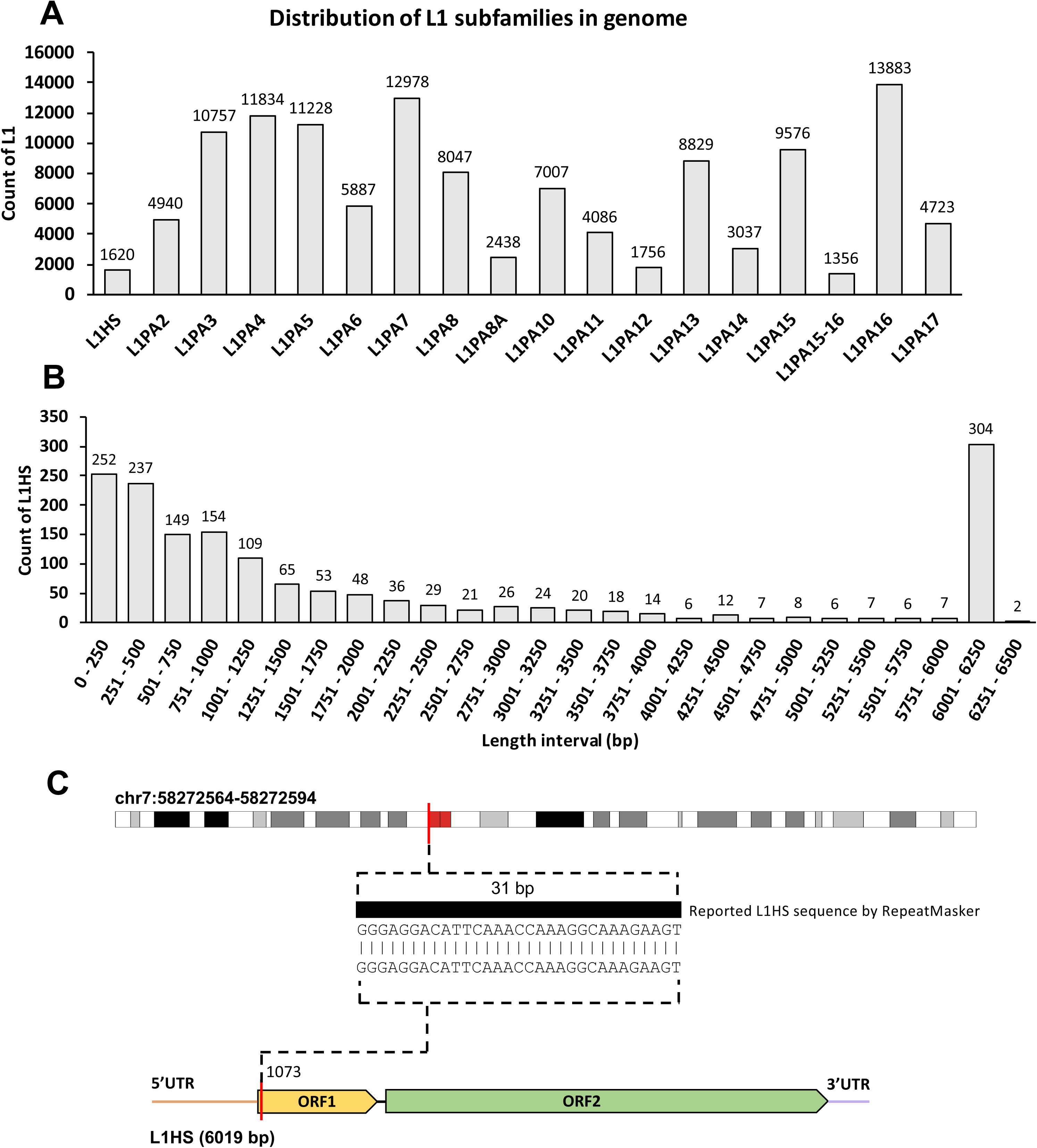
Summarized data of L1 subfamilies reported by RepeatMasker. **A.** Total counts of L1 elements per subfamily. **B.** Distribution of L1HS subfamily elements reported by sequence length intervals. **C.** Sequence of 31 bp reported by RepeatMasker and assigned as a member of the subfamily L1HS. The sequence is on locus chr7: 58272564-58272594 (Hg38) and belongs to an ORF1, position 1073 of an L1HS element.

In this work, we created the web-based resource L1Farm as a result of a comprehensive analysis and annotation of multiple subfamilies of L1 retrotransposons. The databank of L1Farm contains a complete annotation of L1 regions (5’ UTR, ORF1, intron, ORF2 and 3’ UTR) of distinct subfamilies identified in the human genome. We also generated a complete list of annotated genomic *loci* for the full-length and incomplete L1 elements. Next, we also provided a complete annotation of L1 elements in two individual whole assembled genomes at allele-level resolution. Finally, we compare the reference annotation with the individual-based L1 annotations and discuss the potential outcomes of L1Farm.

## Results and Discussion

### A database for annotated L1 retrotransposon elements in humans

Here we present L1Farm, an annotated databank of human L1 retrotransposons. It is composed by 4 annotated datasets: a) regions of L1 (RG-L1); b) full-length L1 (FL-L1); c) regions of L1 at allele-level resolution (AR-L1); d) full-length L1 at allele-level resolution (AF-L1).

To create L1Farm (Fig. 3), we considered the L1 regions of eighteen subfamilies. After the complete analysis considering the human reference Hg38 genome, a total of 322,406 *loci* were detected, with 99% of them corresponding to the 10 youngest L1 subfamilies (L1HS to L1PA10). These subfamilies are known to have higher conservation, at nucleotide level, when compared to older L1 subfamilies (L1PA11 to L1PA16) [4].

**Fig. 3.**
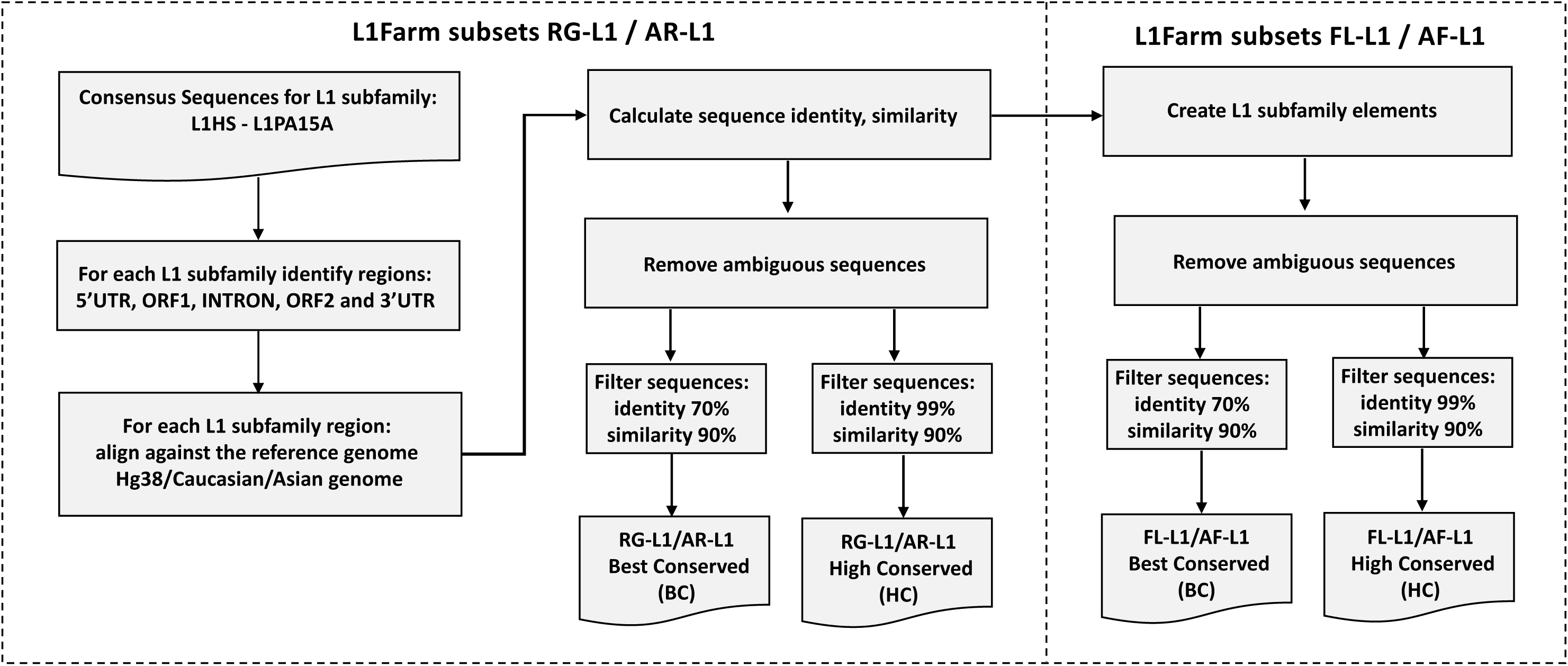
L1Farm flowchart. L1Farm provides the subsets of annotations for L1 regions (RG-L1) and full-length L1 elements (FL-L1), built for the human reference genome (Hg38).In addition, L1Farm explores the distribution of L1 regions (AR-L1) and full-length L1 elements (AF-L1) across individual Caucasian and Asian genomes. All the databases were assembled considering two configurations: minimum sequence similarity of 90% and minimum sequence identity of 70% (BC), and minimum sequence similarity of 90% and minimum sequence identity of 99% (HC).

With the annotated elements, we then investigated the amount of L1 regions through the human reference genome. Using a restricting criterion with a minimum sequence similarity of 90%, we applied two cut-offs: a) 70% (best conserved L1: BC) and b) 99% (highly conserved L1: HC) of sequence identity. L1 regions for the BC and HC have shown variable distribution across all chromosomes, but the absolute amount is directly correlated with the chromosome length, except for the X chromosome that presents more copies of L1s (Fig. S1 and Fig. S2).

For the BC cut-off, the youngest subfamilies from L1HS to L1PA7 correspond to ∼ 90% of the identified elemtns (Fig. 4A-4E and Tables S1-S5). We also found that L1 regions from the investigated subfamilies have higher frequency on the X chromosome, corroborating with previous analyzes performed on human genome sequences [38,39]. Furthermore, the amount of 5’UTR region is less frequent when compared to the other L1 regions (Fig. 4A). This observation is consistent with the statement that the 5’UTR region has the highest variation among L1 lineages [4,5,39].

**Fig. 4.**
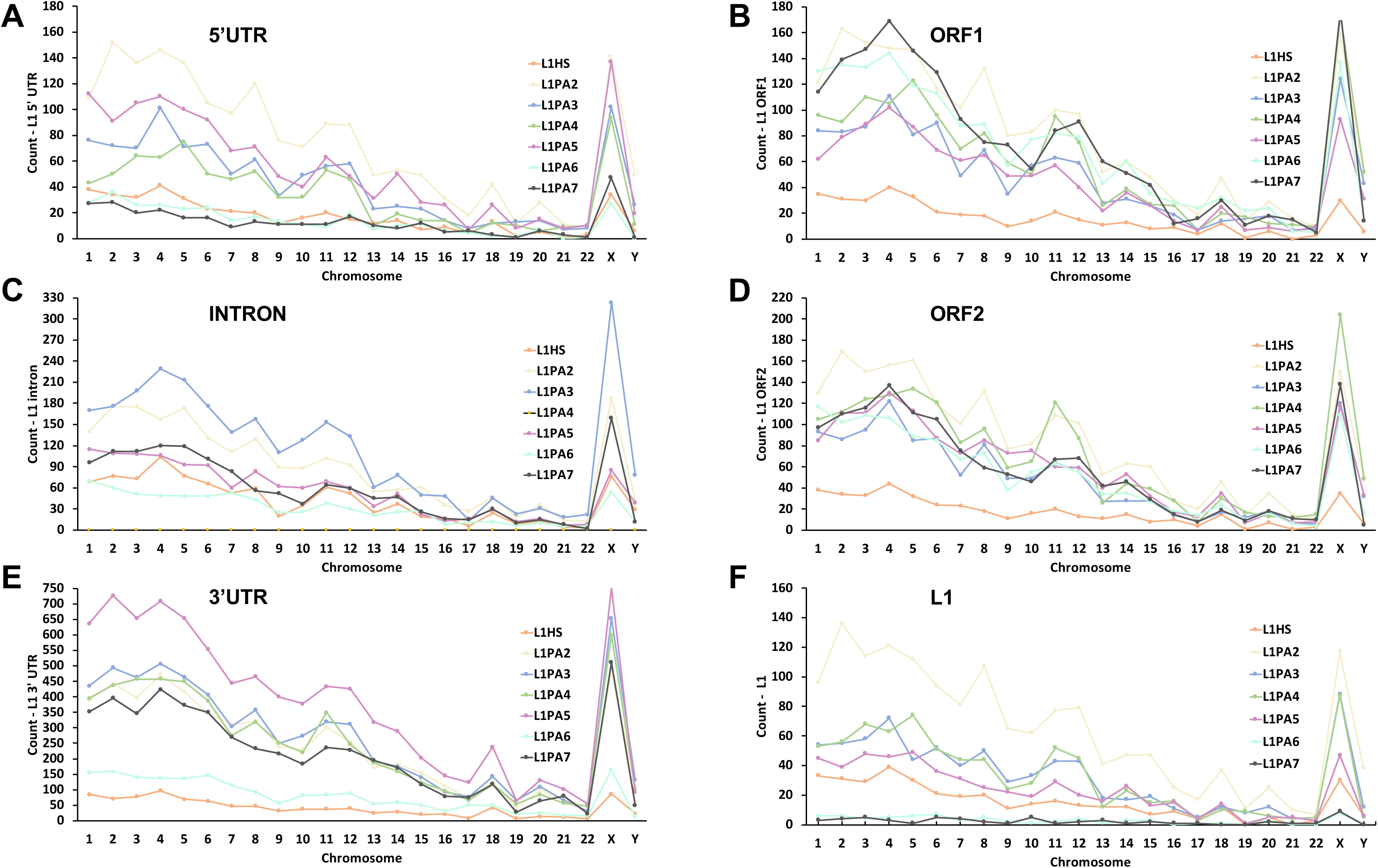
Absolute distribution of L1 subfamilies and regions through human chromosomes. For all comparisons, the subset RG-L1 and the BC cut-off was considered. **A.** Count of 5’UTR regions by L1 subfamily and chromosomes. **B.** Count of ORF1 regions by L1 subfamily and chromosomes. **C.** Count of intron regions by L1 subfamily and chromosomes. **D.** Count of ORF2 regions by L1 subfamily and chromosomes. **E.** Count of 3’UTR regions by L1 subfamily and chromosomes. **F.** Count of full-length L1 subfamilies elements by chromosome.

Using the information of each specific L1 region in the subset of annotations of RG-L1 with the cut-offs BC and HC, we also annotated the genomic locations for those full-length L1 elements using the previous bioinformatics pipeline (Fig. 3).

All the information used to represent the L1 regions (Fig. 4A-4E) and the full-length L1 elements (Fig. 4F) were stored in a subset of L1Farm named FL-L1. This dataset has a complete genome-level panorama of full-length L1 elements in human reference genome (Fig. S3 and Fig. S4), which comes to facilitate future analysis involving L1 subfamilies.

These L1 annotations were all based on Hg38 genome, which is an assembly of multiple sequenced individuals and has multiple variations represented by a single consensus genome. Thus, to complement our understanding of L1 distribution through individual genomes, whether they follow the same characteristics as found in the human reference genome, we took in advantage the availability of the individual whole diploid genomes of two distinct ethnicities, a Caucasian genome and an Asian genome. For these 2 individual genomes, we performed a comprehensive annotation of full-length L1 elements (AF-L1) and its corresponding regions (AR-L1). The annotation process was based on the same protocol (Fig. 3) as the ones applied to the human reference genome, including the assemble of full-length L1 elements using the application FL-Assemble.

Next, we compared all the regions of L1HS subfamily of the Hg38-based RG-L1 subset with the Asian subset AR-L1 and Caucasian subset AR-L1, considering the BC cut-off. For the Asian genome, the distribution of regions through the diploid chromosomes is 98% similar. The same comparison made on Caucasian genome presents less similarity, 83% (Fig. 5). The chromosome 22 presents the main difference when compared to the reference genome: 70% for the Asian genome and 45% for the Caucasian genome (Table S6).

**Fig. 5.**
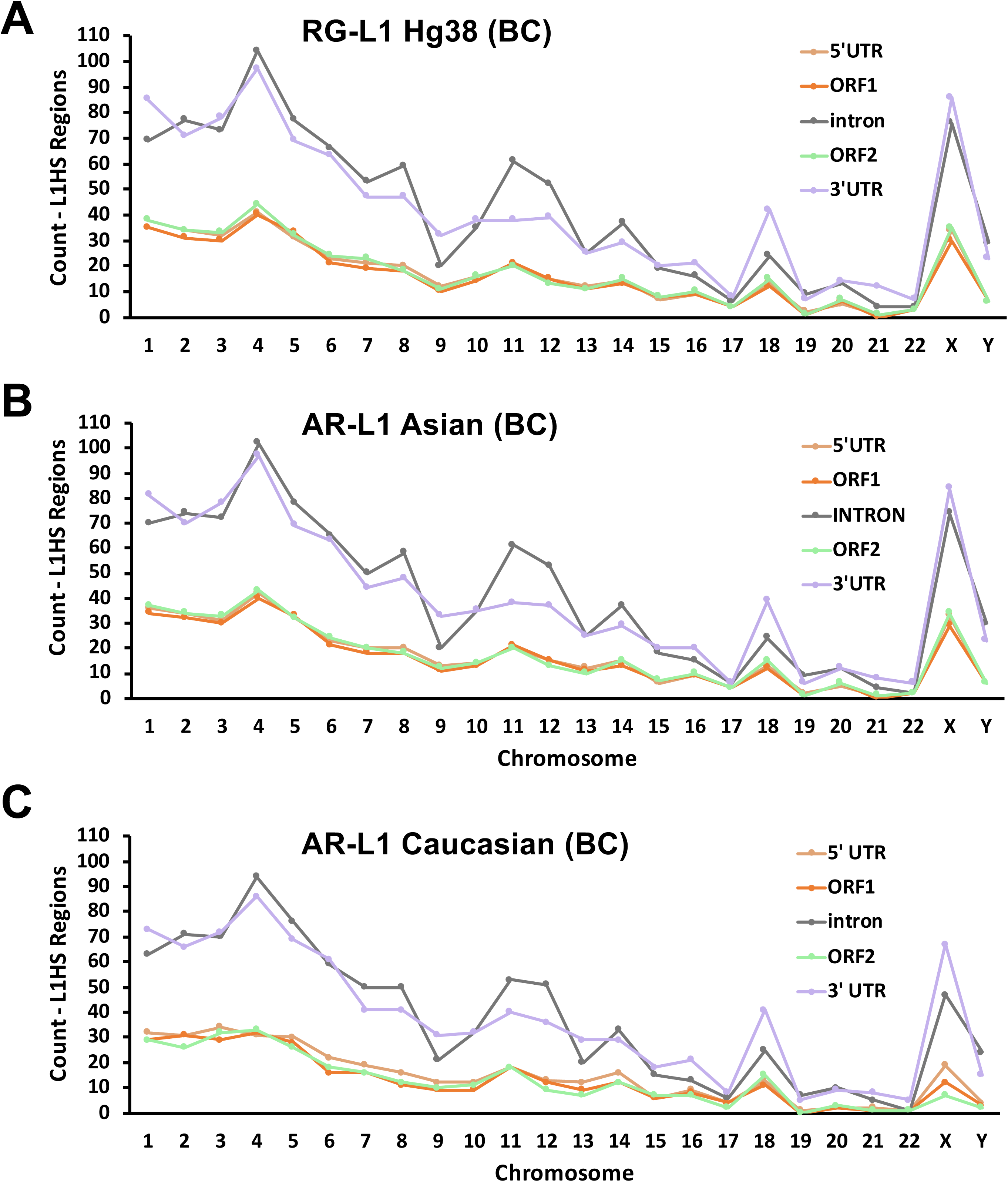
Distribution of regions belonging to the L1HS subfamily among reference genomes. For all comparisons, the BC cut-off was considered. **A.** Distribution of L1HS regions to the subset RG-L1 Hg38 genome. **B.** Distribution of L1HS regions to the subset AR-L1 Asian genome. **C.** Distribution of L1HS regions to the subset AR-L1 Caucasian genome.

We also investigated the distribution of annotated regions of L1HS elements in the subset RG-L1 occurring within protein coding regions (Fig. 6A and Fig. 6B). We found 439 sequences for the BC cut-off and 284 sequences for the HC cut-off, all belonging to L1HS regions. Within these data, several insertions were found in intronic regions of genes that are risk factors for autism such as Stromal Antigen 1 (*STAG1*) [43] and Cadherin 13 (*CDH13*) [44], and intronic regions of genes that are risk factors for cancer such as ATM Serine/Threonine Kinase (*ATM*) [45] and RAD51 Paralog B (*RAD51B*) [46]. Although insertions of L1 elements in intronic regions of genes do not have always a direct impact on the translated protein, they can generate aberrant transcripts that can lead to the development of human diseases [47,48].

**Fig. 6.**
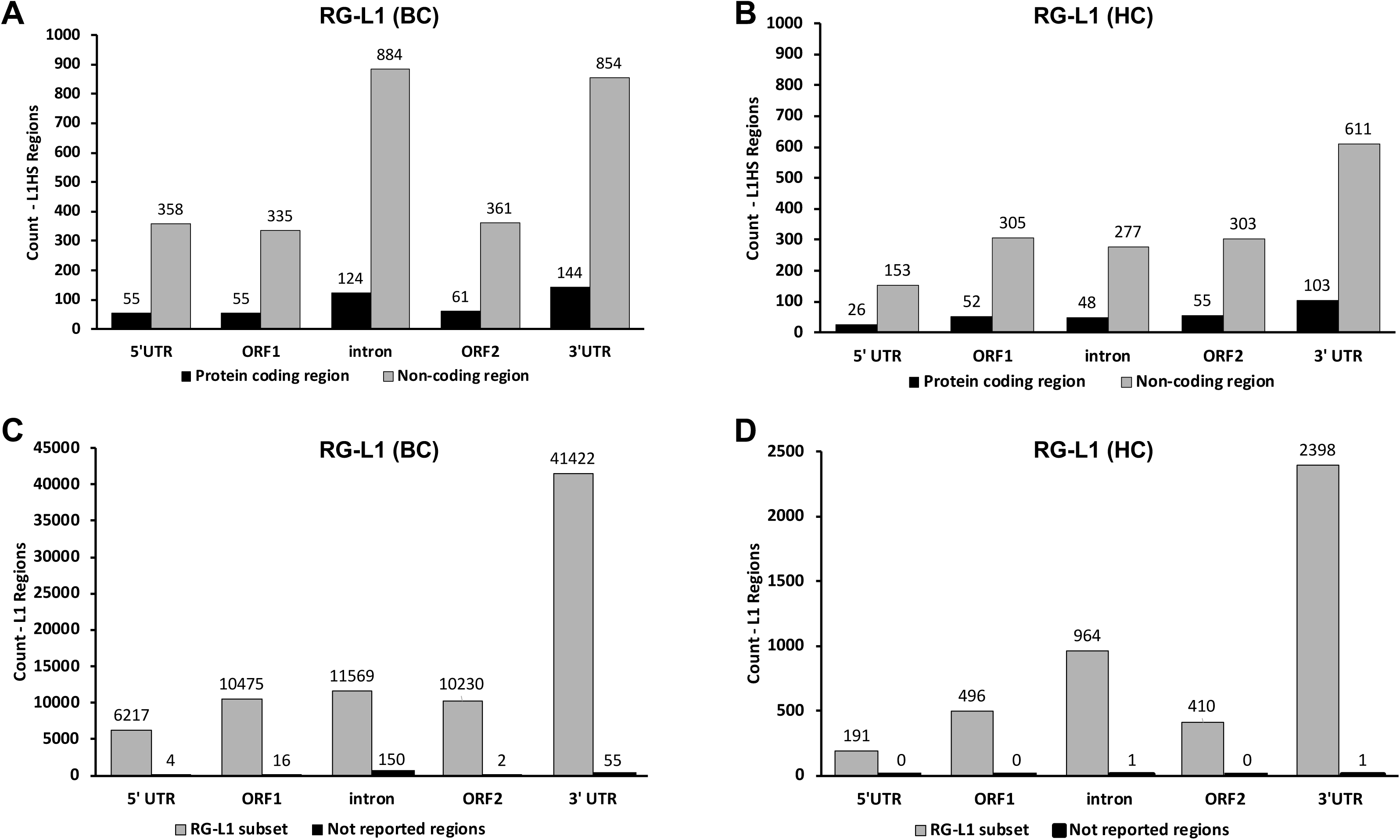
Distribution of the regions of L1HS in protein coding regions and comparison of annotations from RG-L1 subset and RepeatMasker. For all comparisons, the subset of annotations RG-L1 was considered. **A.** Distribution of L1HS regions in protein coding regions with BC cut-off. **B.** Distribution of L1HS regions in protein coding regions with HC cut-off. **C.** Comparison between the L1Farm annotated regions and RepeatMasker with BC cut-off. **D.** Comparison between regions in L1Farm and elements reported by RepeatMasker with HC cut-off.

In addition to the previous analyzes, we also compared the annotations of RG-L1 with the annotated elements reported by RepeatMasker (Fig. 6C and 6D). The criterion for considering that an L1 is in both databases is the existence of at least one overlapping nucleotide and the same sequence strand orientation. Considering the BC cut-off (Fig. 6C), we observed L1 regions annotated in RG-L1 subset that are not reported by RepeatMasker (4 5’ UTR, 16 ORF1, 150 introns, 2 ORF2 and 55 3’ UTR). These 227 unreported elements by RepeatMasker are characterized as belonging to several extinct L1 subfamilies, such as from primates (L1PB) and from mammals (L1MA). One representative example is the 5’ UTR of a L1 within a specific genomic *locus* (Hg38, chrX:152,006,773-152,007,798), which is reported as a sequence that belongs to the subfamily L1PA5 within the RG-L1 subset, that has a sequence identity of 94.2% and similarity of 99.6%, which is described as a L1MA4A element by the RepeatMasker. In another case, a region reported in RG-L1 subset as an ORF2 region (Hg38, chr2:199,073,688-199,077,484) that belongs to the L1PA7 subfamily (sequence identity of 94% and similarity of 99.3%), is redundantly annotated by RepeatMasker as an ancient L1 sequence L1MD2 and as a simple repetitive sequence (AT)n. These conflicting annotations were treated and comprehensively solved L1Farm, through the use of full-length consensus sequences and similarity-based sequence annotations.

## Concluding remarks

Together, the annotated L1 regions (RG-L1, FL-L1, AR-L1 and AF-L1) form the subsets of L1Farm, that provides a catalog of subfamilies of retrotransposons with the best conserved sequences (BC cut-off) and highly conserved sequences (HC cut-off) in humans.

The RG-L1 annotation subset provided by L1Farm differs from others L1 annotations by having elements that contain all L1 regions (regulatory and protein coding regions) and thus offering a panorama of potentially active elements (annotated in the FL-L1 subset). Conflicting repetitive sequence names were treated through the use of full-length consensus sequences and similarity-based sequence annotations.

Furthermore, in order to complement the understanding of L1 distribution through individual genomes, we also included the subset AR-L1 with annotations of L1 regions and the subset AF-L1 with full-length L1s, all at allele-level resolution for the individual Asian and Caucasian genomes.

Thus, L1Farm characterize and annotate L1 elements with detailed information, including similarity-based sequence annotations that can improve *in silico*, *in vitro* and *in vivo* studies to better investigate the role of retrotransposons in human genome.

## Material and Methods

### Retrotransposon reference sequences

The consensus sequences of L1 retrotransposons were obtained from an independent work [4] that created a set of eighteen subfamilies of consensus L1 sequences, from the younger to the older classification: L1HS, L1PA2, L1PA3, L1PA4, L1PA5, L1PA6, L1PA7, L1PA8, L1PA8A, L1PA10, L1PA11, L1PA12, L1PA13B, L1PA13A, L1PA14, L1PA15A, L1PA15B, L1PA16. Next, these consensus L1 elements were fragmented into the following regions: 5’UTR, ORF1, intron, ORF2 and 3’UTR. Although it is a consensus sequence, they have high conservation level at nucleotide resolution. Thus, L1 nucleotide content does not correspond to sequences with multiples distinct variations introduced by individual L1 sequences.

### Sequence alignment

The regions (5’UTR, ORF1, intron, ORF2 and 3’UTR) of the eighteen subfamilies of consensus L1 sequences were aligned to the human reference genome, build Hg38 (assembly GCA_000001405.27), using the Bowtie2 aligner [37] (end-to-end mode, parameters: -a -D 20 -R 3 -N 0 -L 20 -I S,1,0.50).

### Redundancy removal

Alignments were then subjected to the analysis of sequence similarity and sequence identity to treat redundant genomic regions that were associated with regions of two or more distinct L1 subfamilies. Thus, for a genomic region having two or more associated L1 subfamilies, we kept the annotation for the L1 element with the highest identity and similarity, which were later included into the subset of annotations RG-L1.

### Annotation of full-length L1 elements

For this step, we developed an in-house bioinformatics application, named FL-Assemble software, that initially sorts all the annotated regions (5’UTR, ORF1, intron, ORF2 and 3’UTR) for each subfamily according to their genomic positions. Next, FL-Assemble identifies those genomic locations having the five L1 regions in linear order that corresponds to a full-length L1 element, not allowing gap open between the L1 regions that composes a full-length L1 element.

### Personal genome datasets

The personal (individual) genomes used in this study are a Caucasian genome (Craig Venter Individual Genome – assembly GCA_000252825 [40]) and an Asian genome (assembly KOREF_20090224 [41]).

### Protein coding reference annotation

The used protein coding reference annotation was the HAVANA reference database [42], which is a curated annotation for the human reference genome (Hg38). For the annotations, we selected only the well-supported full-length transcripts (HAVANA Transcript Support Level 3,4,5 and NA).

## Abbreviations

AF-L1: Allele-level resolution of L1Farm containing full-length L1s
AR-L1: Allele-level resolution of L1Farm containing regions of L1s
ATM: ATM serine/threonine kinase
BC: Best conserved L1
CDH13: Cadherin 13
FL-Assemble: Application to assemble full-length L1s
FL-L1: L1Farm containing full-length L1s
HC: Highly conserved L1
L1: Long interspersed nuclear element-1
L1Farm: Annotation of L1s in human reference and individual genomes
L1HS: Human-specific L1
LINE-1: Long interspersed nuclear element-1
p53: Tumor Protein P53
RAD51B: RAD51 paralog B
RG-L1: L1Farm containing regions of L1s
RNP: Ribonucleoprotein particle
RUNX3: RUNX family transcription factor 3
STAG1: Stromal antigen 1
TPRT: Target-site primed peverse transcription
TREX1: Three prime repair exonuclease 1
UTR: Untranslated region
SOX2: SRY-box 2
SP1: Sp1 transcription factor
SVA: SINE/VNTR/Alu
ORF: Open reading frame

## Declarations

### Ethics approval and consent to participate

Not applicable.

### Consent for publication

Not applicable.

### Availability of data and material

The databases generated are available at http://bioinfo.deinfo.uepg.br/l1farm.

All files are in tab-separated values (TSV) format.

## Acknowledgements

This study was financed in part by the Coordenação de Aperfeiçoamento de Pessoal de Nível Superior - Brasil (CAPES) - Finance Code 001. AF is recipient of a doctoral fellowship from CAPES/FA (Fundação Araucária). RHH is supported by FA (grant #CP09/2016) and Conselho Nacional de Desenvolvimento Científico e Tecnológico (CNPQ, grant #311438/2022-9 and #402773/2022-5)

## Authors’ contributions

AF: designed and performed the experiments, wrote and revised the manuscript. RHH: designed the experiments, wrote and revised the manuscript. All authors read and approved the final manuscript.

## Declaration of the use of generative AI and AI-assisted technologies in the writing process

The authors declare that they did not use AI-based tools in any part of this manuscript.

